# High Dynamic Range Bacterial Cell Detection in a Novel “Lab-In-A-Comb” for High-Throughput Antibiotic Susceptibility Testing of Clinical Urine Samples

**DOI:** 10.1101/199505

**Authors:** Jeremy Pivetal, Martin J. Woodward, Nuno M. Reis, Alexander D. Edwards

## Abstract

Antibiotic resistance in urinary tract infection is a major global challenge, and improved cost-effective and rapid antibiotic susceptibility tests (AST) are urgently needed to inform correct antibiotic selection. Although microfluidic technology can miniaturise AST, the high dynamic range of pathogen density found in clinical urine samples makes direct testing of clinical samples – rather than testing colonies from overnight agar plates – extremely challenging. We evaluated for the first time how pathogen concentration in urine affects microfluidic AST using a novel microplate-compatible high-throughput microfluidic AST system termed “Lab-on-a-Comb”. When tested with clinical *E. coli* isolates at standardised density, these devices gave identical antibiotic susceptibility profiles to standard disc diffusion and microtitre plate tests. Bacterial detection directly in synthetic urine spiked with clinical *E. coli* UTI isolates was possible over a very large dynamic range of starting cell densities, from 10^3^ – 10^8^ CFU/mL which covers the range of pathogen cell densities found in patient urine. The lowest cell density where cell growth was reproducibly detected optically was 9.6x10^2^ CFU/mL, corresponding to one single CFU detected in a 1 μL microcapillary-an unprecedented level of sensitivity. Cell growth kinetics followed a simple Monod model with fast growth limited by the substrate availability and an estimated doubling time of 24.5 min, indicating optimal *E. coli* growth conditions within these microfluidic devices. There was a trade-off between sensitivity and speed of detection, with 10^5^ CFU/mL detection possible within 2h, but 6h incubation required at 10^3^ CFU/mL.

## INTRODUCTION

Although improved diagnostic technology is widely acknowledged to be essential to combat antimicrobial resistance (AMR), antibiotic susceptibility testing (AST) of clinical samples such as urine remains challenging. Although urinary tract infections (UTI) are often self-limiting, a high prevalence places a significant burden on health systems, and when infections escalate hospital treatment can be required. Current primary care treatment protocols therefore rely on empirical antibiotic administration(1), which exacerbates the clinical and economic burden because of a high level of broad and rapidly changing AMR profile necessitating constant updates to guidelines. Resistance ranges up to ~50% of isolates in OECD countries and as high as >80% outside OECD depending on antibiotic(2) limit the effectiveness of empirical prescribing with consequences for the patient (infection will not be controlled) and the public (emergence and spread of resistance). Even if susceptibility testing is requested, urine culture plus AST takes 2-3 days, during which time infections worsen if the empirical antibiotic is ineffective, or if treatment is delayed prior to culture results.

Several emerging technologies have been developed to the point of large scale uptake into central microbiology laboratories(3, 4). Many labs are now using mass spectroscopy for identification of clinically important microbes by peptide mass fingerprinting to replace traditional analytical microbiology techniques (5). Nucleic acid tests (NAT) such as PCR can rapidly detect specific AMR genes, allowing rapid identification of known resistance, but do not always predict functional antibiotic susceptibility and therefore require constant review and updating to ensure all relevant AMR sequences are screened. Next-generation sequencing does not rely on detection of specific AMR genes, but instead requires advanced bioinformatics analysis which remains prohibitively expensive for mainstream high-volume applications. These technologies still typically require pure colonies from overnight plating, and facilitate centralised identification of known organisms and detection of known strains or resistance genes in fully-equipped labs; however they do not measure the antibiotic susceptibility *phenotype*.

In contrast, *functional* AST, including the current gold standard (6-8) broth microdilution (BMD) performed in a microtitre plate (MTP) and disc diffusion assays performed on agar plates, demonstrate antibiotic suitability directly by identifying phenotypically which antibiotic kills cells or prevents growth. Functional AST currently requires a pure bacterial sample isolated by overnight plating, delaying time-to-result and requiring centralised laboratory equipment. There remains an urgent need for novel AST methods that can directly test patient urine and avoid overnight plating. Ideally new methods would replicate conventional functional assays as closely as possible. For example, miniaturised BMD utilising growth detection similar to current automated lab BMD equipment would accelerate adoption of new technology by permitting simple clinical interpretation of results that would be equivalent to current gold standard AST. Healthcare budgets remain a major constraint, with high prevalence of UTI resulting in very large volume of patients presenting with UTI symptoms in primary care. AMR is more prevalent in parts of the world with most limited healthcare budgets. Very low device costs are therefore required, expensive capital equipment should be avoided, and highly skilled technical sample processing or complex interpretation of results are also undesirable. The high prevalence of UTIs means high sample throughput may also be desirable.

Microfluidic devices offer further miniaturisation beyond MTP wells. Increasing numbers of microfluidic analytical microbiology devices illustrate the great potential for miniaturisation in analytical microbiology (9). An early feasibility study indicated rapid bacterial growth is possible within 200um microchannels, although cell growth was monitored by an impractical method of repeated sample removal that may have aerated the sample, and growth kinetics were not monitored over a broad density range (10). A “millifluidic” system made from simple FEP tubing illustrated that analysis of antimicrobial susceptibility can be miniaturised using microdroplets allowing parallel analysis of large populations of individual cells (11). Whilst this microdroplet system is suited to research applications where highly parallel cell analysis is informative, for testing clinical samples a method for sequentially applying and tracking multiple patient samples is required. Microfluidic devices have also been utilised for quantitative susceptibility testing, for example multi-chamber devices that distribute bacterial sample into chambers pre-loaded with differing antibiotic concentrations (12) or use of microfluidic gradient generation (13). Combining microfluidic culture with digital microscopy and image analysis allows direct observation of the effects of different antibiotics on cell behaviour or growth, allowing rapid AST determination without bulk growth detection (14). An alternative digital readout to optical/microscopic monitoring is electrochemical detection, which has been combined with the conventional colorimetric metabolic dye resazurin to detect bacterial growth in microfluidic devices (15). Most microfluidic microbiology studies have focused either on exploring microdevice engineering science, or on addressing fundamental biology research questions, and commercial application of new technology to clinical sample testing remains a major challenge. One notable exception is a microplate-based microfluidic AST device that has been developed into an fully automated product for testing positive blood cultures to rapidly select antibiotics for sepsis (14, 16). However, this device still requires pre-incubation of blood sample prior to AST testing, in order to grow enough organisms for testing. The droplet-in-tubing millifluidic system has likewise been commercialised offering improved resolution in quantitative laboratory antibiotic susceptibility testing (17) but again pure cultures from overnight plating are required for this detailed lab analysis.

A major technical challenge yet to be addressed in microfluidic systems for clinical microbiology is sample dynamic range. To utilise novel functional antibiotic susceptibility tests for direct urine sample testing – as opposed to testing pure overnight colonies – the devices must be capable of performing across the extremely high dynamic range of pathogen cell density found in infected patients, ranging from below 10^3^ CFU/mL to well over 10^7^ CFU/mL(18). Whilst different clinical diagnostic thresholds of cell densities have been debated for UTI, as many as 30-50% of patients presenting with UTI symptoms have pathogen counts below a traditional cutoff of 10^5^ CFU/mL, and so alternative thresholds as low as 10^2^-10^3^ CFU/mL have been explored. The cost of lowering this diagnostic threshold is increased false positive rate(19). The clinically selected threshold is critically important for microfluidic device design as it dictates the minimum test volume required to be confident of detecting at least one single cell. For example, a 1μL device will have a mean content of only 1 cell/device at a threshold density of 10^3^ CFU/mL. At this mean density, approximately 37% of devices will contain zero CFU, with the absolute number of cells in each device following a Poisson distribution. A lower threshold of 10^2^ CFU/mL would require even larger test volumes. This has two implications for microfluidic functional direct clinical sample AST: firstly device geometry cannot be miniaturised beyond clear sample volume limits, and secondly, the ability to detect growth of very low viable cell numbers – ideally down to a single colony forming unit– is critical.

The challenge is exacerbated by the well-documented impact of cell density on minimum inhibitory concentration across a broad range of standard techniques (20). As well as expanding cell numbers and isolating a pure culture, overnight plating allows adjustment of cell density (typically by estimating cell density by turbidity) prior to AST(7, 8). Varying cell density may also affect the time taken for functional detection of growth vs antibiotic inhibition. Conventional AST assays such as BMD utilise colorimetric or fluorimetric metabolic growth indicators such as resazurin, or monitor turbidity that directly indicates cell density. The rate of substrate conversion or turbidity increase is critically dependent on both cell growth kinetics and starting cell density. Evaluation of the impact of starting bacterial density on cell growth and detection kinetics is therefore critical to microdevice development, and is a major focus of our current study.

Here, we focussed on understanding the impact of sample cell density on microfluidic AST performance. To study the interplay between sample cell density, antibiotic susceptibility, and detection kinetics, very large numbers of microdevices were required. We therefore exploited devices developed previously by our group which combine features of microfluidic devices and dipstick tests (such as uranalysis strips). We recently showed these “Lab-on-a-stick” devices were capable of performing classical functional microbiology tests(21). These devices used a hydrophilic polymer coating to internally modify a fluoropolymer microcapillary film (MCF) that contains a parallel array of ten ~200 micron diameter microcapillaries. MCF is manufactured by melt-extrusion and multiplex test strips can be produced in large batches (22-25), allowing us to make and test hundreds of very low-cost test devices, representing thousands of individual microcapillary devices. Samples are drawn up into the ten capillaries by capillary action, simultaneously releasing and mixing antibiotic with sample. Here, we present a novel high throughput system that exploits large numbers of “Lab-on-a-Stick” devices to perform broth microdilution AST directly in urine samples, in a simple microtitre plate compatible format. The new device, termed a “Lab-on-a-Comb”, was designed to perform as many as 4,800 individual growth assays per 96-well plate.

## Experimental

### Design of 96-well plate compatible “Lab-on-a-Comb” array of microfluidic dipsticks

To study the effect of starting cell density on microfluidic AST and bacterial detection kinetics, and thereby move towards an ultimate objective of developing a microfluidic device for rapid AST directly in UTI patient urine, we needed to analyse growth within large numbers of microdevices in parallel. We therefore developed the Lab-on-a-comb concept (figure 1) which comprises a comblike holder of up to 12 microfluidic test strips with a 9 mm pitch, each formed from a 33mm length of 4.3mm wide 10-capillary MCF manufactured from Teflon-FEP. These test strips were produced rapidly at low cost by pre-treating with polyvinyl alcohol (PVOH) to introduce a hydrophilic coating, and then pre-loading with antibiotic as previously described (ESI figure S1 and Reis et al(21)). Samples combined with growth indicator medium in a conventional MTP can then be tested simply by dipping combs of MCF test strips that draw up 1μL samples by capillary action (figure S1). Each test strip performs 10 parallel analytical microbiology tests, with growth indicated by conversion of a blue metabolic indicator dye to pink then white at high cell density. Since each MCF test strip conducts 10 microfluidic tests with 1μL volume, each comb performs 120 tests per row. By using multiple combs interfaced to a microtiter plate, each 96-well plate can be rapidly screened with thousands of individual 1μL tests with no additional equipment. If 5 combs are dipped per well sampling a volume of ~50uL, 50 tests per well allows 4800 tests per plate (figure S1). This high throughput capacity allowed us to analyse in excess of 1000 1uL microcapillary tests in this initial feasibility study.

**Figure 1.**
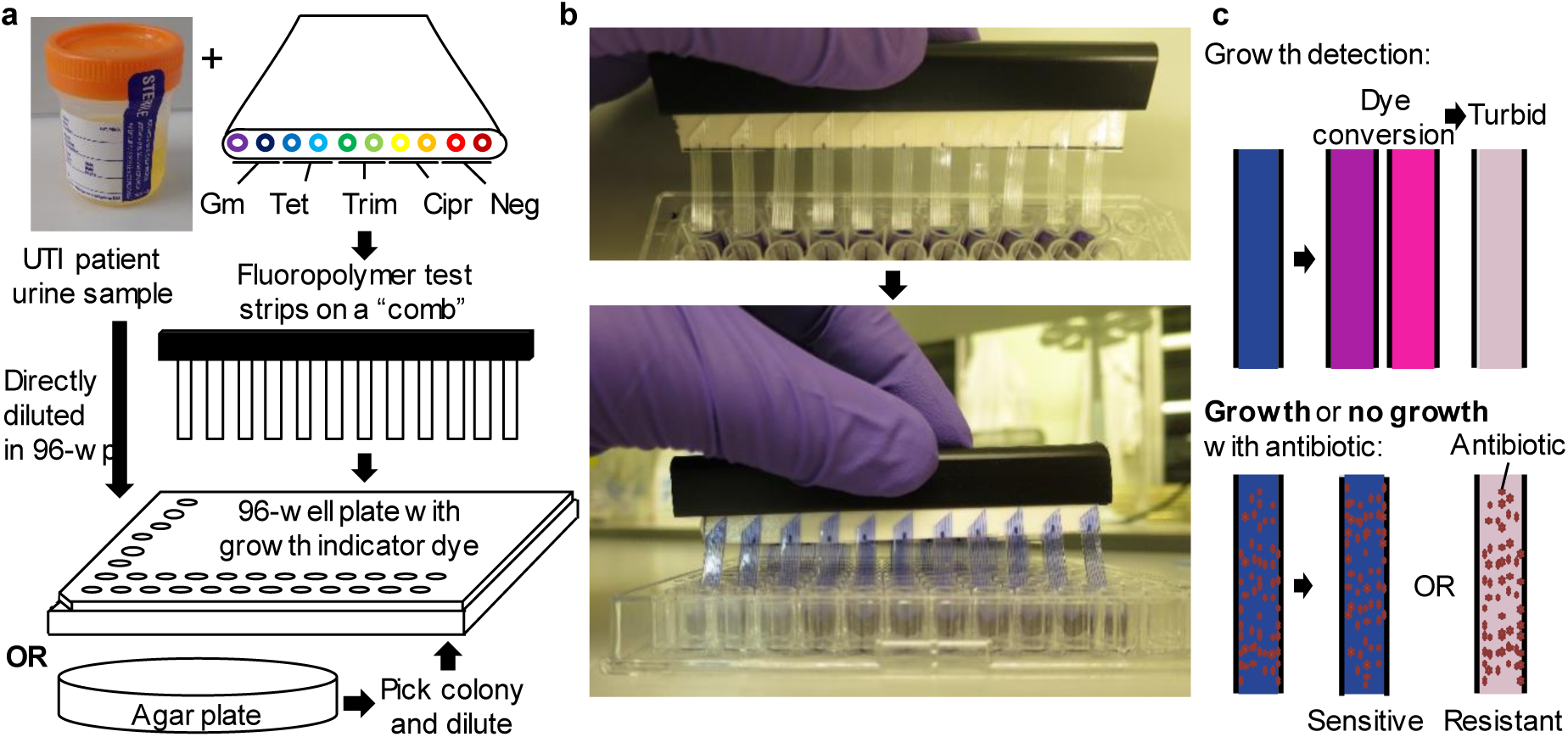
“Lab-on-a-Comb” concept for high throughput, low cost miniaturised functional microbiology. (a) Illustration of UTI urine sample testing protocol using Lab-on-a-Comb device comprising 10-plex microcapillary test strips. (b) Image of Lab-on-a-Comb in use. (c) Cell growth or antibiotic susceptibility detected colorimetrically as resazurin dye changes from blue to pink to white.

### Production of antimicrobial susceptibility dipstick devices

Each comb consisted of a simple plastic paper binder (Office Depot, Andover UK) containing a row of Lab-on-a-Stick test strips made by a method adapted from a previous report demonstrating dipstick microfluidic minimum inhibitory concentration assays (21). The fluorinated ethylene propylene MCF was manufactured by melt-extrusion by Lamina Dielectrics Ltd (Billingshurst, West Sussex, UK) from a highly transparent fluorinated ethylene propylene co-polymer (FEP-Teflon^®^). This material comprises a plastic ribbon containing an array of 10 capillaries along its length with an average diameter of 206 ± 12.6μm and external dimensions of 4.5 ± 0.10 mm wide by 0.6 ± 0.05 mm thick. This material was coated to make internally hydrophilic for production of dipstick microfluidic devices by incubation with a 5 mg/mL solution of PVOH in water (MW 146,000-186,000, >99% hydrolysed, Sigma-Aldrich, UK) at room temperature for a minimum of 2h (26). The material was then washed manually to remove crosslinker with 5 ml of PBS with 0.5 % Tween 20 (Sigma-Aldrich, UK) solution to remove uncoated PVOH, and dried with multiple changes of air using a 50 mL syringe. To pre-load antibiotics onto the internal surface of the PVOH modified MCF, individual microcapillaries of a 1 m long PVOH modified MCF were filled using a 30G needle with freshly prepared filter sterilized antibiotic solutions of gentamicin, tetracycline, trimethoprim and ciprofloxacin (Sigma Aldrich, UK) at concentration of 5 mg/mL in distilled water, 5 mg/mL in distilled water, 3 mg/mL in DMF and 3 mg/mL in 1% HCl respectively. After 5 minutes of incubation, the excess solution was removed by manually injecting air using a 50 mL plastic syringe. All antibiotics were loaded at concentrations that would lead to release of the breakpoint concentration into the sample, taken from published data for *E. coli* (6), based on previously determined loading efficiency quantified by LC-MS(21).

### Bacterial strains

The bacterial strains used in this study were clinical isolates of urinary tract infection strain of *Escherichia coli*, kindly donated by Frimley Park Hospital, UK. Reference strains of *E. coli* ATCC 25922 were used as control for antimicrobial susceptibility testing (LGC group, Middlesex, UK). The bacterial strains were routinely cultivated on Mueller-Hinton agar (Sigma-Aldrich) at 37◦C.

### Dipstick antimicrobial susceptibility testing

Bacterial inoculums were prepared according to BSAC standard for AST (7, 8). Briefly, a well-isolated colony from a fresh (18- to 24-hour culture) bacterial plate were suspended in a saline solution (0.8% NaCl), adjusted to a 0.5 McFarland, diluted 10-fold in a Mueller-Hinton broth (Sigma Aldrich) and further diluted 20-fold in a Mueller-Hinton broth supplemented with 0.25mg/mL of resazurin sodium salt powder (Sigma-Aldrich, UK). The modified antibiotic loaded MCF was trimmed into 40 mm long, dipped into the bacterial inoculum and incubated overnight at 37°C. Test strips were then imaged with a Canon S120 camera using white background illumination to record colour change of the resazurin dye indicator; a change from blue to pink indicating reduction of resazurin and therefore bacterial growth.

To validate the MCF antimicrobial susceptibility method, we performed AST using disc diffusion susceptibility testing following BSAC methods (7, 8). Clearly isolated single colonies from a freshly streaked (18- to 24-hour incubation) bacterial plate were suspended in a saline solution (0.8% NaCl), adjusted to a 0.5 McFarland, diluted 10-fold in a Mueller-Hinton broth (Sigma Aldrich) and further diluted 10-fold in a Mueller-Hinton broth. Inoculum were then spread onto Mueller-Hinton agar plate and antibiotic impregnated discs (Oxoid, UK) of gentamicin (10 μg), tetracycline (30 μg), trimethoprim (2.5 μg) and ciprofloxacin (1 μg) were deposited. Following overnight incubation at 37°C, zones of inhibition surrounding the discs were measured to determine the antibiotic susceptibility profile following the BSAC protocol.

### Evaluating cell growth and detection kinetics across full dynamic range of cell densities

*E. coli* clinical isolates with differing resistance profiles or the sensitive lab strain were serially diluted in synthetic urine to simulate clinical urine samples with a full dynamic range of cell densities. Simulated clinical urine samples were then diluted 5-fold into 96-well plates containing the conventional colorimetric growth indicator dye resazurin at a final concentration of 0.25 mg/mL in Miller Hinton broth. These samples were tested by dipping triplicate Lab-on-a-comb AST devices with duplicate capillaries for no antibiotic vs the 4 common antibiotics tested above, and these devices were then incubated at 37°C. A replicate MTP with identical simulated urine samples diluted in resazurin/broth were incubated alongside the Lab-on-a-Comb devices. Combs and plates were imaged regularly for up to 6 hours with a Canon S120 camera using white background illumination to record colour change of the resazurin indicator dye. The red channel absorbance was calculated at each timepoint for every capillary using ImageJ, and corrected for 0.2mm pathlength as previously described for blue channel absorbance for the immunoassay substrate OPD (23-25). The MTP was imaged and absorbance calculated in parallel using the same camera, white backlight, and image analysis.

### Modelling cell growth kinetics

Bacterial growth in microfluidic strips was modelled following a standard Monod kinetics, by assuming increase in bacteria concentration, *C*_*X*_ and growth rate, *μ* is limited by the concentration of a limiting substrate, *C*_*S*_:

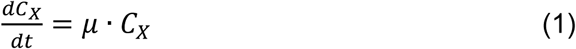

Where

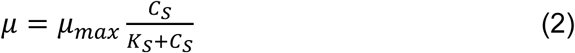

It is possible to define a biomass on subtract yield, *Y*_*X/S*_ as follows:

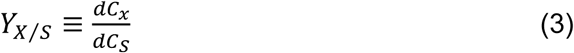

During bacteria growth, the limiting substrate concentration depleted from an initial value, *C*_*S*_(0) to a value at time *t* given by *C*_*S*_(*t*), by analogy biomass concentration increase from initial value, *C*_*X*_(0) to *C*_*X*_(*t*). For simplicity, we assumed subtract concentration correlated with the absorbance value, *Abs*(*t*) measured for each solution. For the microfludic strips, the *Abs* value for each individual capillary were measured from the maximum greyscale peak height from the red channel image split from high resolution (2,400 dpi) RGB images of the microcapillaries. Full details are given in e.g. Barbosa et al. and Castanheira et al.(23-25), and all image analysis was carried out with ImageJ software (NIH, USA). For the case of a 96-well plate, *Abs*(*t*) value was likewise calculated from red channel intensity using ImageJ in wells of digital images of the microplate, shown previously to closely match plate reader absorbance (27), and acquired and analysed in parallel to MCF test strips. For direct comparison of bacteria growth kinetics on the two platforms, all *Abs*(*t*) values herein presented have been converted to units of cm^−1^, by normalising with the *Abs*(*t*) by the maximum light path distance, in this instance 0.02 cm for the microcapillaries and 0.3 cm for the microwells. Although the two optical systems returned distinct *Abs*(*t*) values due to variations in image acquisition, sample geometry and illumination, substrate consumption can be directly compared using the conversion factor, *X*:

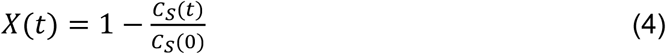

We adapted to *Abs*(*t*) values by assuming at any given time point, *t*:

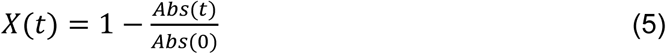

We assumed the initial biomass concentration, *C*_*X*_(0) is related to CFU/ml of the inoculum/sample:

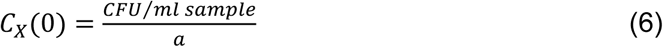

where *a* is an arbitrary parameter with units (CFU mg^−1^). Integration of Monod model in Eqs. (1)-(2) using partial fractions yields the time *t* required to drop *Abs* signal from its initial value, *Abs*(*0*) to a given value *Abs* (all in cm^−1^).

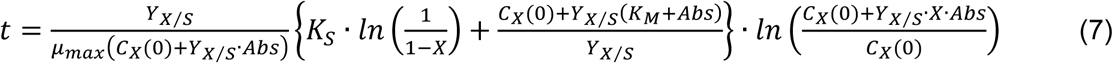

The unknown parameters in the model were maximum specific growth rate, *μ*_*max*_ (h^−1^), the half-velocity constant, *k*_*M*_ (in units of absorbance), and the yield of biomass on substrate, *Y*_*X/S*_ (dimensionless), and the arbitrary parameter *a* (CFU mg^−1^). The model in Aq. (7) was best fitted to the experimental data using Excel’s solver using as criteria minimum square differences. The iterative process resulted in a number of possible solutions to the equation, so we used experimental data to determine the product *Y*_*X/S*_\**μ*_*max*_. It can be shown that during exponential stage of growth, the slope from a In(*Abs*(*t*)) versus *t* plot yields *Y*_*X/S*_\**μ*_*max*_ (again assuming *Abs* value correlated with substrate concentration). Consequently, we have averaged all slopes at different initial CFUs/ml values to yield an average *Y*_*X/S*_\**μ*_*max*_ that was fed into the kinetic model given by Eq. (7). We repeated the procedure for every single individual microcapillary and microwell, yielding the model parameters herein show in Results and Discussion section.

## RESULTS AND DISCUSSION

### Lab-on-a-comb end-point antibiograms deliver accurate susceptibility profiles similar to disc diffusion and microplate methods

To validate the Lab-on-a-Comb concept we adapted the BSAC standard threshold AST (6) from microtitre plates into MCF test strips for four common antibiotics. Disc diffusion in agar plates and endpoint resazurin AST in Lab-on-a-Comb devices were performed on 9 clinical UTI isolates plus a reference *E. coli* sample (ATCC strain 25922). To eliminate the impact of bacterial cell density on growth detection in the presence of antibiotic, a conventional preparation method was followed whereby a fresh colony was suspended in saline with turbidity adjusted to 0.5 MacFarlane, then diluted 100-fold in appropriate indicator medium. The only difference was that MCF has a ~20-fold shorter pathlength than MTP wells and therefore the concentration of resazurin indicator dye was increased to ensure colour change was clearly visible. Endpoint growth was recorded after overnight growth, with white turbid capillaries indicating growth thus antibiotic resistance, and susceptible samples remained blue (figure 2). We found full agreement between the novel and gold standard methods, with identical antibiotic resistance profile of all samples between disc diffusion and Lab-on-a-Comb (Table 1). Whilst overnight plating and cell density adjustment is still required prior to testing, this confirms that Lab-on-a-Comb is an extremely simple but potentially extremely powerful device to increase AST throughput using very low cost microdevices whilst retaining conventional MTP lab format for sample preparation and directly replicating gold standard assay methodology.

**Figure 2.**
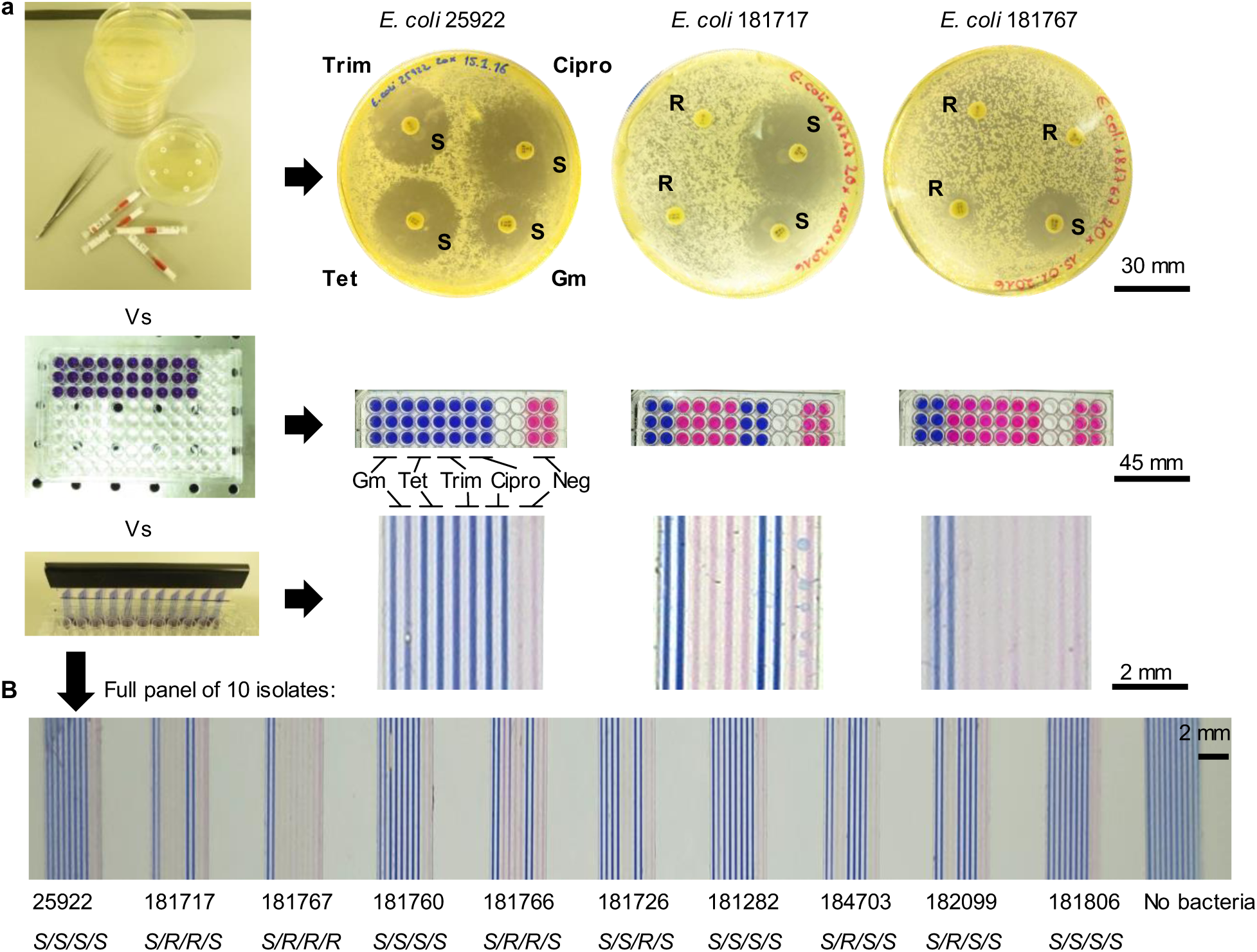
Comparison of miniaturised “Lab-on-a-Comb” method with conventional BSAC disc diffusion and microplate methods for testing the susceptibility of *E. coli* to gentamicin, tetracycline, trimethoprim, and ciprofloxacin with a panel of 9 clinical urinary tract infection isolates plus a susceptible reference strain. (a) illustration of three parallel methods and (b) images of 11 MCF test strips after dipping in the panel of 10 samples and overnight incubation, showing expected antibiotic susceptibility profiles from the conventionally scored profiles as indicated underneath (s=susceptible, r=resistant).

**TABLE 1.** Susceptibility of clinical *E.coli* isolates to four antibiotics

| *Escherichia coli*<br>isolate | Antibiotics |  |  |  |
| --- | --- | --- | --- | --- |
|  | Gm | Tet | Trim | Cipr |
| 1- ATCC 25922 | S | S | S | S |
| 2- 181717 | S | R | R | S |
| 3- 181767 | S | R | R | R |
| 4- 181760 | S | S | S | S |
| 5- 181766 | S | R | R | S |
| 6- 181726 | S | S | R | S |
| 7- 181282 | S | S | S | S |
| 8- 184703 | S | R | S | S |
| 9- 182099 | S | R | S | S |
| 10- 181806 | S | S | S | S |

Gm, gentamicin; Tet, tetracycline; Trim, trimethoprim; Cipr, ciprofloxacin; S, sensitive; R; resistant

### Reproducible growth detection across wide dynamic range of starting cell density

To explore the feasibility of direct testing and study the impact of variable pathogen density in urine, a panel of simulated patient samples was prepared in MTP comprising synthetic urine spiked with 5-fold dilutions of representative *E. coli* clinical isolates demonstrating resistance and susceptibility to our antibiotic panel. This panel was tested with Lab-on-a-Comb antibiogram devices and colourimetric growth detection recorded at regular intervals over a 6h incubation. As expected, blue to pink to white transition was fastest at high cell densities. When analysed, the biggest change in images was a reduction in red light absorbance. This reflects the major peak shift to shorter wavelengths for the absorbance of resazurin following conversion to its product resorufin. This peak shift results in a significant drop in absorbance in the red channel of the digital RGB sensor, which accounts for the apparent colour change from blue (strong red and green absorbance leaving only blue light) to cyan (i.e. pink, with only green channel absorbance leaving red and blue light). The drop in red absorbance was highly reproducible, with duplicate capillaries in triplicate samples showing very consistent kinetics of colour change (figures 3 and S2). Each 33mm long strip contains ten 0.1mm radius microcapillaires each with a volume of 1.0mm^3^, and it was informative to consider growth kinetics vs cell density expressed per 1μl device. Growth was detected within 2.5hr for densities above 6x10^5^ CFU/mL, 4-5h for cell densities 10^4^ – 10^5^ CFU/mL, and 5-6h for density between 10^3^ to 4x10^3^ CFU/mL.

**Figure 3.**
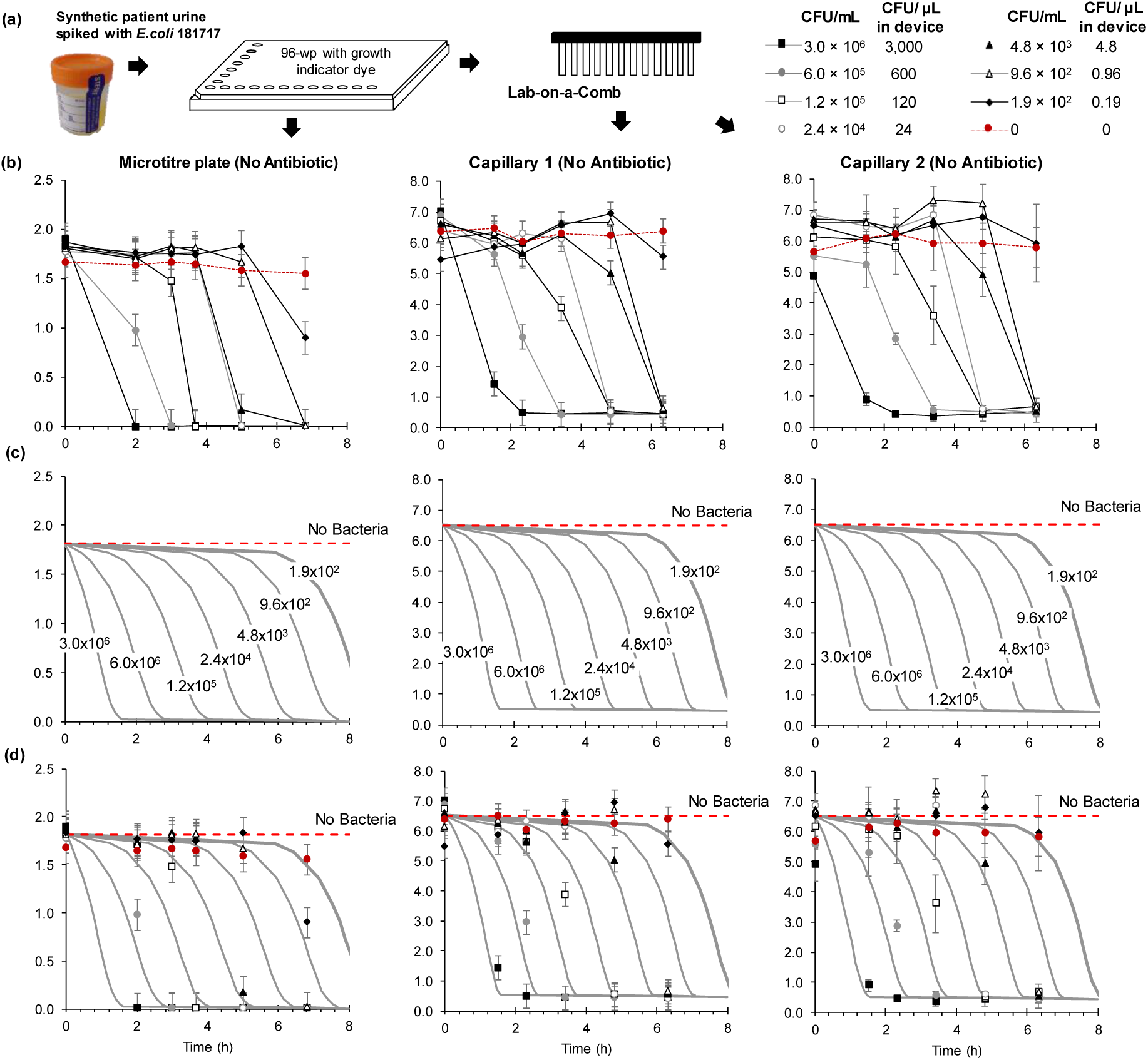
Comparison of detection kinetics from simulated clinical urine samples between microplate and replicate Lab-on-a-Comb microcapillaries at clinically relevant range of starting pathogen density. Clinical isolate 181717 was diluted into simulated urine at the indicated concentrations and resazurin growth indicator medium added in microwell plates. One plate was then tested in three replicate Lab-on-a-Stick test strips loaded in duplicate with no antibiotic in capillary 1 and 2, (with duplicates of 4 antibiotics in capillaries 3-10), and incubated at 37°C alongside a second plate which was incubated directly in the MTP. Plates and test strips were imaged at the indicated times, and red absorbance measured to indicate conversion of resazurin by bacterial growth. (a) Schematic of sample processing into microtitre plate and subsequently in “Lab-on-a-comb” devices. (b) Experimental data showing capillaries 1 and 2 from triplicate test strips, with the mean absorbance plotted +/- 1 standard deviation. (c) Modelling growth kinetics. (d) Combined experimental and numerical results, showing the quality of fit.

Notably, growth was detected at a limiting concentration where approximately 1 CFU per device was expected. Poisson statistics would suggest as many as 36% of capillaries would have no CFU at this cell density, but 6 out of 6 capillaries (3x replica strips each with 2x replica capillaries) in this study were positive and none negative. No positive conversion was seen in MCF at 5x and 25x lower dilutions, although growth was detected as low as 1.9x10^2^ CFU/mL in the larger sample volume of the MTP (figure 3b and data not shown). Given the fast growth rate of these samples, it is possible that cell density at sampling was in fact slightly higher than that determined by plating samples taken of the dilutions and culturing overnight. Overnight plating can also underestimate the number of viable organisms capable of growth in liquid culture.

To detect concentrations lower than 10^3^ CFU/mL in clinical situations, where as few as 10^2^ CFU/mL could indicate infection (19), a larger sample volume can be easily achieved, either using a longer test strip or larger diameter microcapillary, or simply by using multiple replicate microcapillaries-each single MCF strip has a total volume of 10μL therefore 10^2^ CFU/mL corresponds to approx. 1 CFU per strip.

The problem of low cell density being challenging and slow to detect in microdevices has been previously addressed by use of microfluidic cell capture followed by electrochemical growth detection(15), although this introduced a need to pre-filter urine samples to avoid clogging by particulates in urine. However, in spite of enrichment, this device still required sample cell density of 10^5^ CFU/mL for rapid detection, i.e. 100 CFU per μL which is 100x higher than the clinical threshold for UTI diagnosis. Our current devices, whilst slower, are therefore 100x more sensitive than the microcapture device disclosed in Besant et al.(15).

When compared to bacteria detection in MTP wells, the detection of growth was remarkably consistent (figure 3b), which is not surprising as metabolic dye conversion is not diffusion-limited, and therefore one of the major speed benefits of assay miniaturisation is not expected for this mode of bacterial growth detection.

### Kinetics of detection reveals rapid bacterial growth in FEP microdevices

Functional AST inherently requires detection of bacterial cell growth, therefore a detailed understanding of growth kinetics of bacterial cells in microfluidic devices is essential. For rapid detection of low concentrations of cells in clinical diagnostic applications, it is critical to understand if growth in microfluidic devices occurs as rapidly as in conventional culture, and how growth rate is affected by cell density over the clinically important – and very large - dynamic range. To our knowledge this has not previously been reported.

When the kinetics of resazurin conversion was monitored within individual capillaries, the time taken for detectable colour change (loss of red absorbance) increased as starting cell density decreased (figure 3b), suggesting detection follows a conventional bacteria growth model. Cell growth rates were initially estimated from experimental data shown in figure 3b for MCF and MTP wells by determining the mean increase in time to detectable colour change for each 5-fold dilution of starting cell density, measured as 0.92h (MCF) and 0.94h (MTP), which corresponds to a doubling time of 24 minutes.

In addition, we noticed cell growth kinetics followed a Monod model with substrate limitation, which yielded a mean doubling time, *t*_*1/2*_ of 24.5 minutes, as summarised in Table 2, which approaches the expected minimum doubling time for optimum exponential growth of *E. coli* of ~20 minutes. Figures 3c-d directly compares the Monod kinetic model in a MTP well and two separate MCF capillaries from triplicate test strips, with the best-fitted parameters summarised in Table 2. Bacteria growth followed same growth model across experimental replicas and the two different platforms, sharing same values for kinetic parameters *μ*_*max*_ and *a*. The best-fitted values of *Y*_*X/S*_ and *K*_*M*_ were however distinct between the MCF capillaries and MTP wells due to the different optical setups. Our model as shown in Eq. (7) is linked to absorbance values (cm^−1^), therefore dependent on the extinction coefficient for the optical setup, however allows direct comparison between different platforms. Two main observations resulted from best-fitting of kinetic model to experimental data: firstly, experimental replicas in MCF capillaries perfectly match, and secondly bacteria growth in microcapillaries follows same kinetics established for macrofluidic systems.

**TABLE 2.** Kinetic model parameters

|  | MICROTITER<br>PLATE | CAPILLARY 1 | CAPILLARY 2 |
| --- | --- | --- | --- |
| Biomass on substrate yield, $Y_{X/S}$ [-] | 1.82 | 0.85 | 0.83 |
| Substrate 'half velocity' constant, $k_m$ [abs units] | 0.5 | 1.0 | 1.0 |
| Maximum growth rate, $\mu_{max}$ [$h^{-1}$] | | 1.7 | |
| Arbitrary parameter for conversion CFU/ml to<br>biomass concentration, $a$ [cfu $mg^{-1}$] | | $3.0 \times 10^6$ | |
| Doubling time, $t_{1/2}$ [min] | | 24.5 | |

### Detection of antibiotic susceptibility by growth inhibition

Whilst the detection of low cell densities was promising, and the standardised cell dilutions showed identical susceptibility profiles to conventional AST, it was important to understand how cell density might affect antibiotic susceptibility detection. The strain tested was resistant to tetracycline and trimethoprim (figure 2), and growth kinetics were identical to no antibiotic, as expected (figure 4). The kinetics of growth detection was highly reproducible between duplicate capillaries loaded with antibiotic (figure S3). At the lower starting cell densities, the expected growth inhibition by antibiotics gentamicin and ciprofloxacin was also observed (figure 4). However, a threshold density was clear for some antibiotics, above which growth detected in the presence of antibiotics. Starting cell density is known to affect *in vitro* growth inhibition assays (20). But even when colour change was observed in the presence ciprofloxacin (at the highest cell density tested), the rate of bacteria growth was dramatically slowed compared to no antibiotic, and the red channel absorbance fell less and never reached 1.0, as seen with all starting cell densities without antibiotic or with resistance; this suggests susceptibility may be detectable through differential kinetics, rather than by endpoint bulk colour change. In contrast, with gentamicin, the red channel absorbance eventually fell to 1.0 for the two highest cell densities, although 2-4h later than without antibiotic. Interestingly, the growth inhibition in parallel MTP well samples was more clear-cut than in microcapillaries (figure S4). There are two possible reasons for this difference. Firstly, our antibiotic loading and release method used in the “Lab-on-a-Stick” devices initially delivers a gradient of reagent along the length of the capillaries (21), with higher concentration at the top meniscus. Whilst this can eventually equilibrate by molecular diffusion of reagents during incubation, at high cell densities resazurin conversion may occur quickly enough for colour change before the antibiotic concentration becomes homogeneous along the full length of the microcapillary. Modified reagent loading and release methods are being explored to overcome this effect, which can also be minimised by reducing the length of the test strip. Secondly, it is possible that inhibition of metabolic activity is different between the different platforms, presenting somehow distinct microenvironments. We are looking to adapt in the future the kinetic growth model shown in Eq. (7) to consider antibiotic inhibition and couple it with a multiscale model that will allow determination of microenvironmental differences. Additional data is required to inform if microfluidic AST testing requires selection of different threshold antibiotic concentrations compared to MTP wells, however our proof-of-concept data in figure 4 fully demonstrates the feasibility of miniaturising AST.

**Figure 4.**
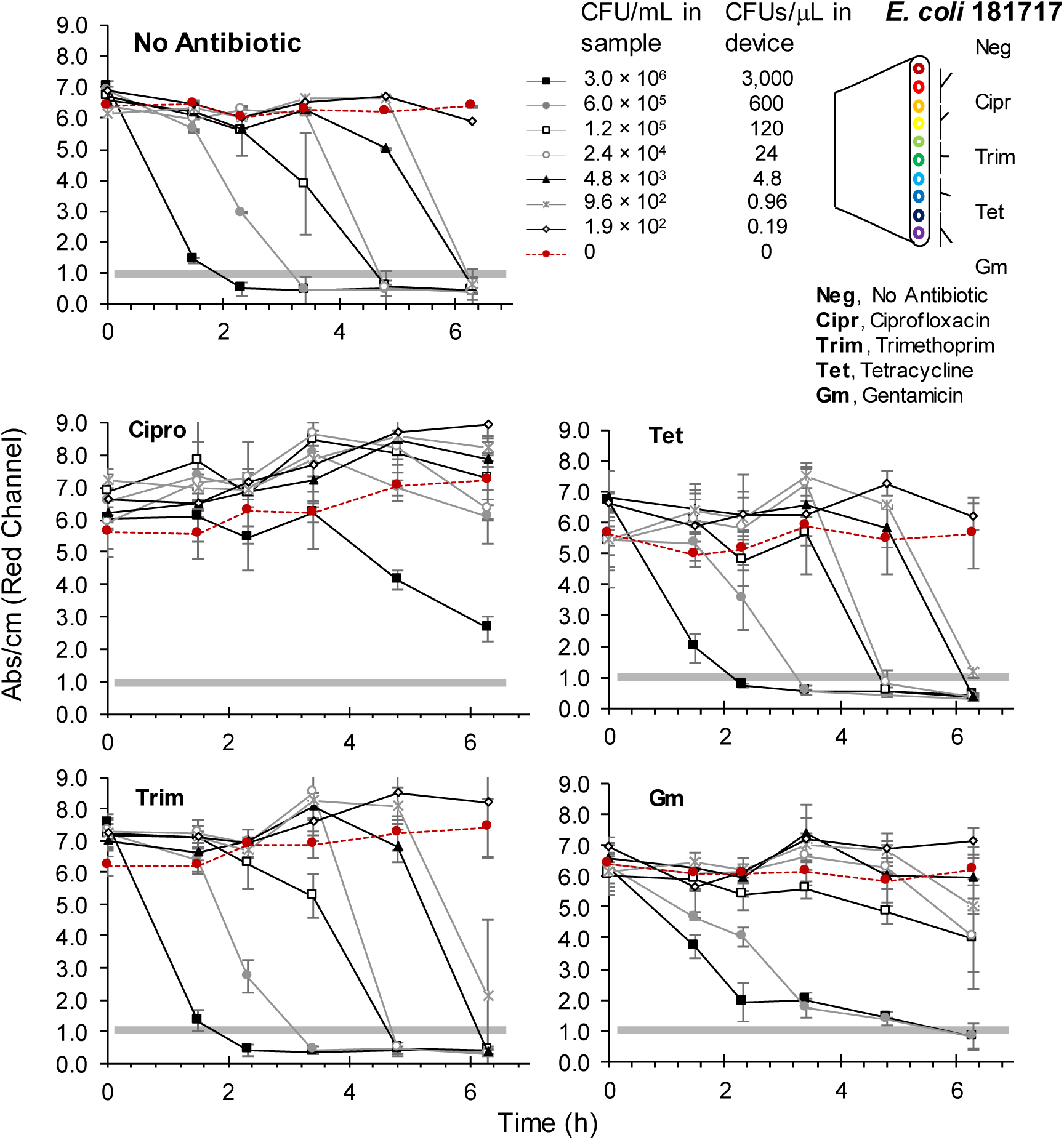
Impact of starting cell density on kinetics of metabolic growth detection in microcapillaries in the presence of antibiotics. As indicated in figure 3, *E. coli* isolate 181717 was resuspended in simulated urine at the indicated concentrations and tested in Lab-on-a-Stick devices with duplicate capillaries containing the indicated 4 antibiotics, or no antibiotic. Capillaries 1, 3, 5, 7 and 9 of test strips are shown, with the mean red absorbance of triplicate test strips plotted with error bars indicating +/- 1 standard deviation. The second replicated capillary showed the same kinetics of resazurin conversion.

### Feasibility of direct AST in clinical urine samples

This preliminary evaluation demonstrates the potential of MTP-compatible low-cost microfluidic devices for performing AST in a clinical microbiology lab using conventional functional assays. Whilst a broad range of cell density could be detected in simulated patient samples, suggesting a step towards near-patient diagnostics, further development is still required to achieve direct urine testing. Firstly, even faster detection of growth is needed, with colorimetric resazurin detection known to be hard to colorimetrically quantify by digital imaging due to complex changes in visible light absorbance (27). Faster detection may be achieved by switching to fluorimetric growth detection, previously used by our group for immunoassays in MCF (26). Secondly, a better understanding of the predictive value of direct urine AST testing (without overnight plating and colony isolation) in clinical samples is vital. An area for future work is the heterogeneity of clinical urine samples, with variable levels of contamination from commensal organisms.-Other new technologies have been explored for direct sample testing, for example microscopic growth analysis of samples in MTPs explored the feasibility of direct AST in clinical samples (rather than pure isolates), albeit in infected urine from catheterised pigs (28).

## Conclusions

Using our new “Lab-on-a-Comb” high-throughput device we reported for the first time a detailed analysis in microdevices of how the kinetics of detection and functional antibiotic susceptibility testing of the UTI pathogen *E. coli* is affected by sample cell density. We demonstrate that cell growth vs inhibition can be distinguished in simulated infected urine samples across the >4 log_10_ dynamic range relevant to clinical urine samples. The time-to-result depends strongly on cell density, and ranged from 2-6hr, nevertheless bacteria growth in microcapillaries closely followed that observed for the macrofluidic MTP well. Growth was consistently detected within a single microcapillary with cell densities as low as 9.6x10^2^ CFU/mL, suggesting growth of a single CFU can be reliably detected with “Lab-on-a-Comb” in a maximum of 6 hours. However, an impact of starting cell density on detection of antibiotic susceptibility was observed, with the highest starting cell densities showing some false positive growth in the presence of some antibiotics, incorrectly indicating antibiotic resistance in a susceptible organism. The extent of this problem depended on the antibiotic. This suggests that some optimisation of threshold antibiotic concentration may be required in miniaturised devices, to account for this variability in growth behaviour in different assay formats. This initial study provides new understanding of the consequences of miniaturisation on vital functional analytical microbiology assays, whilst taking a step towards two critically important applications: increasing throughput of laboratory AST methods, and ultimately towards near-patient rapid urine testing to inform antibiotic selection for treating UTI.

## Figure Legends

**Figure S1**

Further details of “Lab-on-a-Comb” concept.

**Figure S2**

Replicates of bacterial growth detection kinetics in multiple capillaries and multiple isolates.

**Figure S3**

Replicates of bacterial growth inhibition in the presence of antibiotics in multiple capillaries.

**Figure S4**

Kinetics of bacterial growth inhibition measured in microtitre plate wells.

